# Experimental production of anti-membrane bound variable surface glycoglycoprotein (anti-VSGm) antibodies for *in vitro* and *in situ* immunostaining of pathogenic African trypanosomes

**DOI:** 10.1101/2020.03.13.990465

**Authors:** Reymick Okwong-Oketch, Julius Nsubuga, Peter Ayebare, Zachary Nsadha, George William Lubega, Ann Nanteza

**Affiliations:** College of Humanities and Socio-sciences, Makerere University, Kampala, Uganda; College of Veterinary Medicine, Animal Resources and Biosecurity, Makerere University, Kampala, Uganda

**Keywords:** Trypanosome, variable surface glycoprotein, immunostaining, antibodies, bloodstream

## Abstract

**Background:** The variant surface glycoprotein (VSG) of the African trypanosomes is the major membrane protein of the plasma membrane of the bloodstream stage of the parasite. African trypanosomiasis (sleeping sickness in humans and nagana in animals) is caused by the systemic infection of the host by several sub-species of the extracellular haemoflagellate protozoa under genus Trypanosoma. As a defense barrier against the host immune response, the entire surface of the bloodstream form of trypanosome is covered with densely packed molecules of VSG that determines the antigenic phenotype of the parasite. Variant surface glycoprotein has a C-terminal domain that is highly conserved in various species of trypanosomes.

**Methods:** The membrane bound VSG (VSGm) protein was prepared without denaturing the homologous region and by including numerous variable antigen types from *Trypanosoma brucei brucei* parasites. The purified VSGm native trypanosome protein was used to produce anti-VSGm immune sera in rabbits. The indirect immunofluorescence assay (IFA) was used to detect trypanosomes from mice blood, artificial culture media and cattle histological sections.

**Results:** The resultant immune sera were able to detect different strains and species of African trypanosomes from *in vivo* and *in situ* sources after immunostaining. Anti-VSGm antibodies also demonstrated a unique property to locate trypanosomes within the histological tissues even after the trypanosome’s morphology had been distorted.

**Conclusion:** The produced immune sera can be utilized for immunohistochemistry to detect *Trypanosoma species* in various fluids and tissues.

**Author summary:** 

## Introduction

African trypanosomiasis results from systemic infection of the mammalian host by several sub-species of the extracellular hemoflagellate of the genus Trypanosoma. The parasites are mainly transmitted by a bite of the blood-sucking tsetse fly of the genus *Glossina* (1), and after the initial skin infection, they multiply extracellularly (2) by binary fission in the bloodstream, lymph and interstitial fluids (3), then spread via the lymph and blood circulation within the host (4). Human African trypanosomiasis (HAT) in divided into an early hemolymphatic stage, in which the parasites are confined to the blood and lymph, and a late encephalitic stage, in which *T. brucei* and/or increased protein content and numbers of lymphocytes are found in the cerebrospinal fluid (CSF) (5).

The drugs widely used in the treatment of second-stage HAT are highly toxic (6,7), therefore, both diagnosis and staging of the disease must be highly accurate as a correct diagnosis is beneficial for both infected individuals and the community (8). Diagnosis of trypanosomes follows a three-step pathway (screening, diagnostic confirmation and staging). Diagnostic confirmation mostly relies on identification of trypanosomes in the blood, lymph nodes or CSF. However, the standard parasitological techniques available miss up to 30% of patients (9). Staging allows classification of HAT into the hemolymphatic or meningo-encephalitic stage but because of the absence of reliable blood tests able to detect central nervous system (CNS) invasion by the parasite, staging relies on the CSF examination (8). According to (10) there is no consensus as to what exactly defines a biologically CNS disease or what specific CSF findings should justify drug therapy for late-stage involvement yet diagnosis of the CNS involvement is crucial in order to give an appropriate treatment regimen.

Trypanosomes have a complex antigenic structure and elicit the production of a large spectrum of antibodies. Variant surface glycoprotein (VSG) is specific to trypanosomes thus anti-VSG antibodies that are able to stain various strains and/or species of trypanosomes would be very significant in mass screening and staging of human trypanosomiasis. Such antibodies would for example be coated on plates and be used for mass screening, detecting recently released VSG or trypanosomes in blood. This could reduce on incidences of false positives and false negatives common in the card antigen test for *Trypanosoma brucei gambiense* (CATT/*T. b. gambiense*) (11), but also reduce on time lag between sampling and examination to avoid immobilization and subsequent lysis of trypanosomes in the samples.

Microscopic examination of lymph node aspirate, blood or CSF provides direct evidence for trypanosome infections. However, most of the available parasite detection methods would not detect the parasites if their number is below the detection limit of 100 trypanosomes/ml (12). Anti-VSG antibodies, however, can detect even small amounts of the VSG protein in released or soluble form, thus could be used to supplement the current methods used in microscopic examination, especially of CSF obtained by lumbar puncture during disease staging in HAT. This could allow detecting VSG protein in CSF even after ten minutes of sample collection in which trypanosomes begin to lyse (8), but also promote sensitivity of trypanosome detection in CSF at less time, unlike the double centrifugation method which is time consuming and requires two centrifuges (8,13). This study produced anti-VSG immune sera in rabbits that could be used to stain heterologous strains and species of African trypanosomes *in vitro* and *in situ.*

## Methods

### Ethics statement and considerations

The study was carried out in accordance with the recommendations in the Standard Operating Procedures (SOPs) of experimental animal care of the College of Veterinary Medicine, Animal Resources and Biosecurity (COVAB), Makerere University and adequate consideration of the 3R’s (Replacement of animal with non-animal techniques, Reduction in the number of animals used and Refinement of techniques and procedures that reduce pain and distress). The protocols for animal management, handling and blood collection were approved by COVAB Institutional Animal Care and Use Ethical Committee (Reference Numb; SBLS.AN.2017).

### Experimental animals and trypanosome stocks

All the experimental animals were purchased from the COVAB, Small Animal Laboratory Experimental Unit, Makerere University, Kampala, Uganda. Twelve (12) weeks old Albino rats and four weeks old Albino mice were used to propagate trypanosomes. Fourteen (14) weeks old New Zealand White rabbits were used to produce anti-VSG immune sera following their immunization by the purified VSGm native trypanosome protein. All control animals were sex and age-matched with test animals in the immunization protocol. Trypanosome stabilates made from *T. b. brucei UTRO 010291B*, obtained from Molecular Biology Laboratory, Makerere University, Kampala, Uganda, courtesy of Prof. Enock Matovu were used for parasite propagation leading to VSG protein extraction. Frozen stabilates had been prepared as described by (14).

### Propagation of *Trypanosoma brucei brucei UTRO 010291B* in mice

All experimental infections for parasite propagations were made after at least one mouse passage from frozen stabilate to revitalize the trypanosomes. Mice were always infected by intraperitoneal injections of 1 × 10^4^ trypanosomes in 200µl of Phosphate Saline Glucose (PSG) buffer. The trypanosomes were propagated in three (3) mice and parasitaemia monitored daily by wet preparations from tail blood and its levels determined by Matching method (15).

### Purification of *Trypanosoma brucei brucei parasites* from whole blood

This was done by chromatography using diethylaminoethyl (DEAE)-cellulose (Sigma-Aldrich, Germany); regenerated according to manufacturer’s protocol and packed into a column (50ml syringe) using the gravity flow, then equilibrated using PSG buffer until the effluent was pH 8. The rats were sacrificed when the parasitemia was high (10^8.2^-10^8.4^ trypanosomes/ml of blood) and blood was drawn by cardiac puncture into 8ml EDTA vacutainer tubes. The EDTA blood was loaded into the column and the components allowed to flow into the DEAE cellulose. Then 1x PSG (pH 8) buffer was continuously added to the column until all the trypanosomes had passed through it as determined by checking the effluent microscopically at intervals. An aliquot (10µl) was used to count the number of trypanosomes in the effluent using a Neubauer haemocytometer. The effluent was then centrifuged at 3500 revolutions per minute (rpm) for twenty minutes, the supernatant was poured off and the mass/weight of the pellet determined. The pellets were snap frozen and stored in liquid nitrogen until required for protein extraction.

### Purification of the *Trypanosome brucei brucei* VSGm protein

This was performed as described by (16). Briefly, the *T. b. brucei* bloodstream form cell pellets (approximately 5×10^10^ cells) was retrieved from −80°C storage and re-suspended in 20ml of Krebs-Ringer phosphate buffer (128mM NaCl, 5mM KCl, 2.8mM, CaCl_2_, 1.2mM MgSO4, 10mM glucose, 10mM Phosphate buffer (pH 7.4). The suspension was treated on ice with an equal volume of 10% (w/v) Trichloroacetic acid and the precipitate centrifuged at 9000xg for 10 seconds to obtain a loose pellet. It was re-suspended in 4ml of distilled deionized water. The suspension was extracted with 20 volumes of chloroform/methanol (2:1) (v/v) with vigorous shaking for 5 minutes and stored overnight at 4°C to increase the protein yield. The extract was then separated into 2 phases by addition of 0.2 volumes of 0.9% NaCl solution and centrifuged at 12000xg for 1 hour at 4°C. The upper aqueous phase containing purified VSGm was recovered by aspiration and dialyzed against 3 × 20 litres of distilled de-ionized water at 4°C for 36 hours. The retained material was freeze-dried and resultant VSGm stored at 4° C. Prior to use, the solid VSGm protein was reconstituted in 1ml of 1x PBS. An aliquot (10µl) was used for protein quantification and analysis. The bulk of reconstituted VSGm was stored in 0.2ml aliquots at −80°C.

### Detection of *T. b. brucei* VSGm protein by SDS-PAGE analysis

Sodium dodecyl sulfate-polyacrylamide gel electrophoresis (SDS-PAGE) was performed (17,18) followed by staining of proteins with Coomassie brilliant blue.

### Quantification of *T. b. brucei* VSGm protein

The Bradford assay (19) for the quantification of proteins was used. Ten standard solutions (1ml each) containing 0.05, 0.10, 0.15, 0.20, 0.25, 0.30, 0.35, 0.40, 0.45 and 0.50mg/ml Bovine Serum Albumin (BSA) (Sigma-Aldrich, Germany) in 1x PBS were prepared. The control (10µl) of 1x PBS alone, then the 10 aliquots of BSA standard solutions were loaded into the wells in the microtiter plate, 10µl of VSGm to be quantified was also loaded in triplicate (dilutions of 1:1, 1:2, 1:4, and so on in 1x PBS were made basing on the protein concentrations). Then 200µl of the Bradford reagent (Sigma-Aldrich, Germany) diluted five-fold in distilled water was loaded into all the microplate wells containing the samples and the plate was incubated for 10 minutes at room temperature with gentle shaking to mix the samples in the wells. The absorbance of the blue colour of the Bradford dye by the samples was measured at a wavelength of 595nm in a spectrophotometer (Biotek, EL 800) and a standard curve was drawn from the optical density values to obtain the concentration of VSGm.

### Production of *T. b. brucei* anti-VSGm antibodies in rabbits

#### Immunization of rabbits

Each test rabbit was immunized with 200µg of purified VSGm protein re-suspended in 0.5ml of 1x PBS and emulsified with 0.5ml of Qual A adjuvant. Each control rabbit was injected with 0.5ml of Qual A adjuvant alone in 0.5ml of 1x PBS buffer. The rabbits were injected subcutaneously at four different sites with 0.25ml per site of the uniform emulsion. Boosting was done twice at 14 day intervals from the day of immunization, each time using 50µg of VSGm re-suspended in 0.5ml of Qual A for each test rabbit, and 0.5ml of Qual A adjuvant alone in 0.5ml of 1x PBS buffer for each control rabbit. The experiment was run in duplicates to determine the consistency of the results.

#### Collection and processing of rabbit sera

Blood (10ml/rabbit) for pre-immune sera preparation were obtained from the rabbits two days before immunization, and blood for immune sera was obtained on days 12 and 26 post-immunizations. The recovered sera (approximately 5ml/rabbit) were kept at −20°C until use.

#### Analysis of rabbit sera by Western blotting

The VSGm protein samples were separated on 10% SDS-PAGE then trans-blotted onto two separate nitrocellulose membranes using the mini PROTEAN II Trans-blot unit (Biorad). The membranes were rinsed in 1x PBS (0.1% w/v) Tween 20 and fixed with 0.2% glutaraldehyde (ICN Biomedicals) to improve on the retention of antibodies. The blots were blocked for 1 hour with 1x PBS containing 0.5% BSA to reduce non-specific protein binding sites. One membrane was incubated with anti-VSG immune serum as primary antibody (diluted 1:1,000 in 0.5% (w/v) skimmed milk in 1x PBS containing 0.05% BSA and Tween 20 (0.1% w/v). The other membrane was incubated in pre-immune serum at the same dilution. Membranes were then probed with horse radish peroxidase (HRP) conjugated goat anti-rabbit IgG (Sigma-Aldrich, Germany) (diluted 1:10000 in 0.5% w/v skimmed milk in 1x PBS containing 0.05% BSA and Tween 20 (0.1% w/v) as secondary antibody. Then membranes were incubated in 1.3mM Diaminobenzidine (DAB) solubilized in 1x PBS containing 0.04% hydrogen peroxide (H_2_O_2_) as the substrate to visualize any positive reactions. The reaction was then stopped by addition of double distilled water (ddH_2_O). Sera at different experimental stages from the test and control rabbits were tested during the analysis. Incubations with primary and secondary antibodies were carried out for one hour per incubation under gentle shaking. Washing of the membranes after each incubation was done 3 times for 5 minutes per wash using 1x PBS/Tween 20 (0.1% w/v).

### Immunofluorescence assay (IFA) on *T. b. brucei UTRO 010291B* from various sources

The IFA for VSGm staining of bloodstream form *T. b. brucei* UTRO 010291B parasites Trypanosomes from *in vivo* (mice) or *in vitro* (artificial media cultures) (provided by other studies) were incubated with pre-immune or immune sera (for the two immunization experiments) and then probed with anti-rabbit IgG conjugated to FITC. The IFA for VSGm staining of bloodstream form *T. b. brucei* UTRO 010291B and other species and strains was also carried out. Briefly, the viability of trypanosomes was determined microscopically by motility. About 4 × 10^7^ trypanosomes in 10µl of blood were fixed on each of the poly-L-lysine pre-coated glass slides and incubated at room temperature for 20 minutes to allow trypanosomes to stick onto the glass slides. Trypanosomes on separate slides were incubated in a humidified chamber at room temperature with anti-VSGm antibodies or pre-immune serum each prepared at a dilution of 1:4000 in PBST. After an hour, the primary antibody was discarded and the preparations probed with a secondary antibody (anti-rabbit, conjugated to Fluorescein isothiocyanate, FITC (Sigma-Aldrich, Germany), prepared at a dilution of 1:10000 in PBST for 1 hour. The slides were washed three times in 1x PBST and air dried at room temperature in the dark. The cells were then viewed under a fluorescent miroscope with an oil immersion objective (X 100). The IFA for VSGm protein staining of other various bloodstream form trypanosome species and strains included *T. b. brucei GVR35, T. b. brucei 427, T. b. rhodensiense UGA 15, T. b. rhodensiense UGA 17* and *T. congolense BA 59).* These parasites were also retrieved from liquid nitrogen as stabilates and propagated in mice prior analysis to ensure their viability by motility after revitalization.

### The IFA for VSGm staining of procyclic form *T. b. brucei UTRO 010291B* parasites

This procedure was similar as for the bloodstream form trypanosomes above, with the exception of procyclic form trypanosomes propagated in artificial culture media used instead of bloodstream form trypanosomes. The parasites were cultured in a CO_2_ incubator at 37°C.

### Immunostaining of *T. b. brucei* parasites in infected cattle histological brain sections

The procedure was done on archieved cattle brain tissue histological sections fixed on microscopic slides. The slides were provided from another study where cattle were experimentally infected to study the pathogenesis of *T. b. brucei* in cattle (Courtesy of Prof. GW Lubega). A total of 27 slides were used; 18 slides (test slides; from infected cattle) had duplicate brain sections that had showed positive results with trypanosomes with hematoxylin and eosin (H&E) staining. The other nine slides (the controls; from non-infected cattle) had showed negative results with H&E staining. The brain sections were dewaxed by dipping the slides in two changes of 100% xylene, each for five minutes and then dehydrated by dipping in two changes of 100% ethanol, each for three minutes, 95% and 80% ethanol, each for one minute and then rinsed in distilled deionized water. Antigen retrieval was then performed by dipping the slides in pre-heated Sodium citrate buffer (10mM Sodium citrate, 0.05% Tween 20, pH 6) for 25 minutes. The slides were cooled at room temperature and rinsed twice in PBST, two minutes/rinse. They were blocked with 5% BSA in PBST (0.1% Tween 20) for 30 minutes at room temperature. The sections were then incubated with primary antibodies (immune and pre-immune sera) prepared at a dilution of 1:4000 in PBST. After an hour, the primary antibodies were discarded and the sections probed with secondary antibody (anti-rabbit IgG conjugated to FITC) (Sigma-Aldrich, Germany) prepared at a dilution of 1:10000 in PBST. The slides were then washed 3 times in 1x PBST and dried at room temperature in the dark. The sections were viewed under a fluorescent microscope (Leica, DFC310 FX) with an oil immersion objective (x 100) in the dark room.

## Results

### The recovery of purified *T. b. brucei VSGm* protein

The VSGm prepared was of the expected molecular weight (61kDa) as shown in Fig. 1

**Figure 1:**
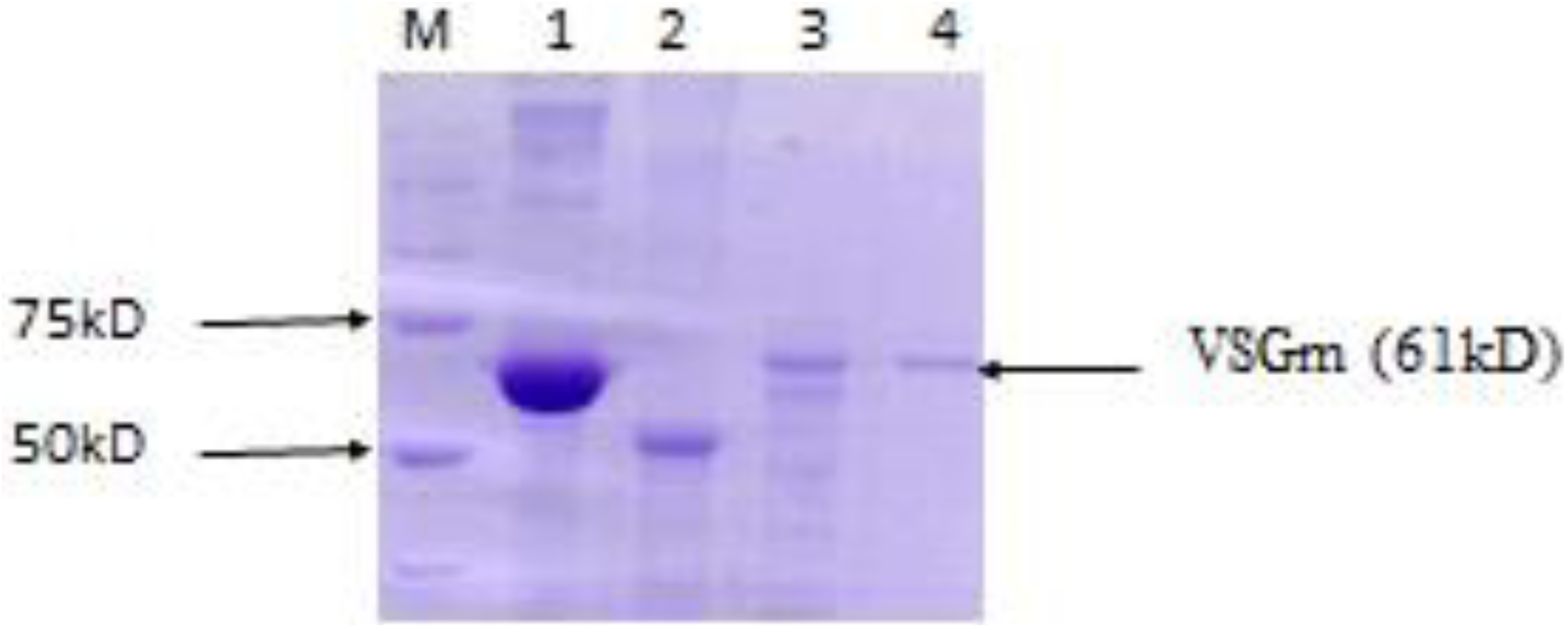
Polyacrylamide-gel electrophoresis analysis of the *T. b. brucei* VSGm protein. The 10% SDS-PAGE gel was used to detect the purified VSGm protein. Lane M: Precision plus protein standard marker, lane 1: Bovine serum albumin (BSA), lane 2: Recombinant trypanosome β-tubulin, lanes 3 and 4: *Trypanosoma brucei* VSGm proteins (prepared as separate batches). The purified *T. brucei* VSGm was of the expected molecular weight (61kDa).

### The specificity of *T. b. brucei* anti-VSGm antibodies

The anti-VSGm antibodies produced in rabbits were highly specific to VSGm protein as they stained only *T. b. brucei* VSGm but not bovine serum albumin (BSA) (a mammalian protein) or trypanosome β tubulin on Western blotting (Fig. 2).

**Figure 2:**
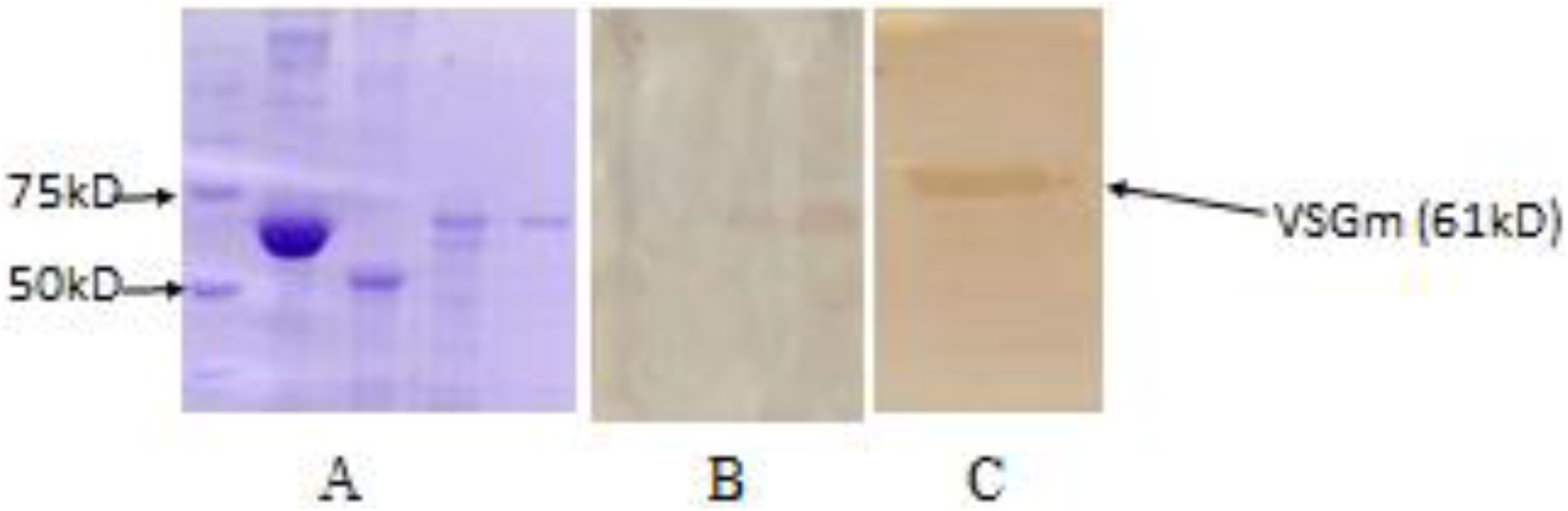
The SDS-PAGE and Western blot analysis of the rabbit *T. b. brucei* anti-VSGm antibodies. The VSGm were electrophoresed on a 10% SDS-PAGE alongside BSA and recombinant trypanosome β tubulin and stained with Coomassie Brilliant blue (Panel A) or transferred to nitrocellulose paper and probed with immune serum (Panel B or C). Lane M: Precision plus protein standard marker (Sigma), Lane 1: BSA, Lane 2: Recombinant trypanosome β-tubulin protein, lanes 3 and 4: *Trypanosoma b. brucei* VSGm. The immune sera were specific as could only detect th*e T.brucei* VSGm proteins (Panels B & C; lanes 3 and 4).

### Trypanosome detection by Immunofluorescence assay

#### The bloodstream form trypanosomes immunofluorescence analysis

The bloodsteam form *T. b. brucei* UTRO 010291B, non-permeabilized trypanosomes probed with anti-VSGm antibodies (dilution 1:4000) showed clear signals/staining, with the staining uniformly distributed over the surface of the cells. Non-permeabilized trypanosomes probed with monoclonal anti-β-tubulin (dilution 1:4000), pre-immune sera (dilution 1:4000) and PBS did not show any signals/staining. Permeabilized trypanosomes probed with anti-β-tubulin (dilution 1:200) gave clear signals/staining of the cells. Non-permeabilized trypanosomes probed with anti-β-tubulin at a dilution 1:200 did not show any signal/staining (Fig. 3).

**Fig. 3:**
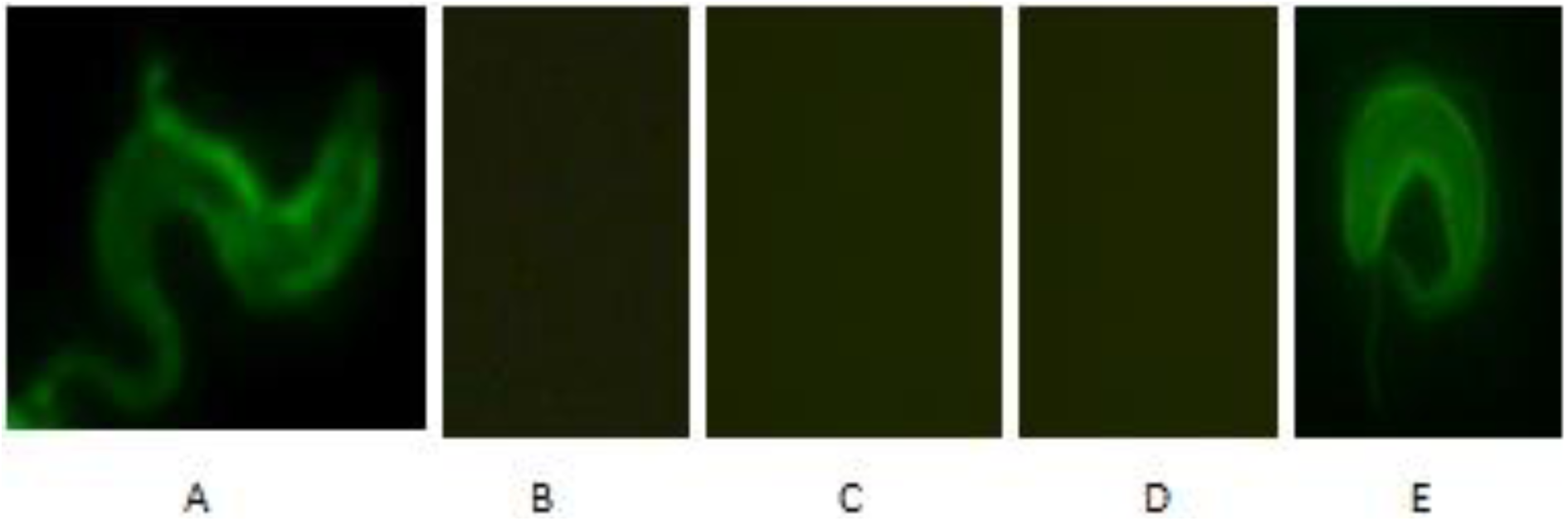
Immunofluorescence of bloodstream forms of *T. b. brucei* from *in vivo* culture. Bloodstream form (BSF) trypanosomes were cultured and processed (without permeabilization) for immunofluorescence analysis with anti-VSGm immune serum (panel A), pre-immune serum (panel B), 1x PBS (panel C) or anti-trypanosome β-tubulin (panel D) at dilutions of 1:4000. Permeabilized BSF trypanosomes probed with anti-trypanosome β-tubulin at a dilution of 1:200 (panel E).

#### The procyclic form trypanosomes immunofluorescence analysis

The *in vitro* propagated procyclic form *T. b. brucei* UTRO 010291B showed clear fluorescence signals with permeabilized trypanosomes probed with anti-β-tubulin (dilution 1:200) (positive control) but not with non-permeabilized trypanosomes probed with anti-VSGm immune serum, pre-immune serum, anti-β-tubulin or PBS (Fig. 4).

**Fig. 4:**
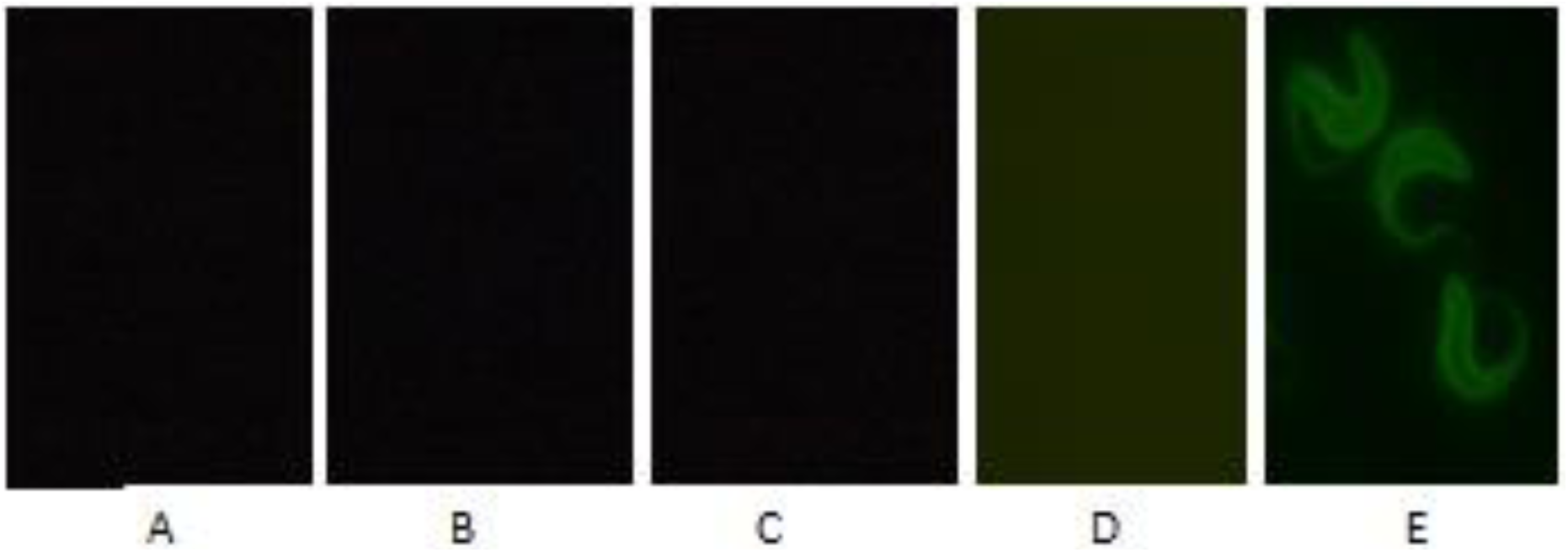
Immunofluorescence analysis of procyclic form *T. b. brucei* from *in vitro* culture. Procyclic form trypanosomes (*T. b. brucei*) were cultured and processed (without permeabilization) for immunofluorescence analysis with anti-VSGm immune serum (panel A), pre-immune serum (panel B), 1x PBS (panel C) or anti-trypanosome β-tubulin (panel D) at dilutions of 1:4000 as described in the methods. Panel E: permeabilized procyclic form trypanosomes probed with anti-trypanosome β-tubulin at a dilution of 1:200. The anti-VSGm antibodies could not detect the procyclic form trypanosomes.

#### The heterologous trypanosome species and strains VSGm immunofluorescence analysis

Clear signals were also obtained with the heterologous non-permeabilized bloodstream form *T. b. brucei* GVR35, *T. b. brucei* 427, *T. b. rhodensiense UGA 15, T. b. rhodensiense UGA 17* and *T. congolense BA 59*, as shown in Fig. 5 below.

**Fig. 5:**
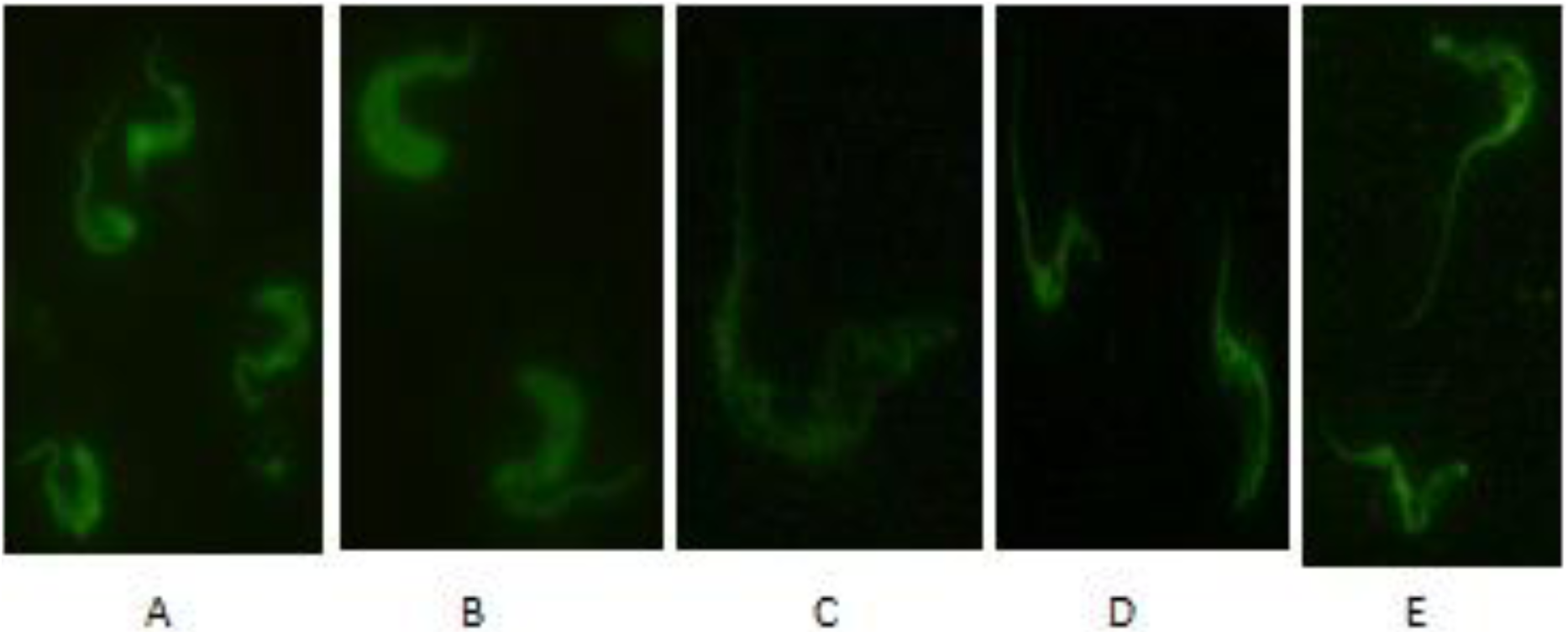
Immunofluorescence staining of heterologous bloodstream form trypanosome species and strains by anti-*T. b. brucei* VSGm antibodies. Bloodstream forms of *T. b. brucei* 427 (Panel A), *T. congolense BA59* (Panel B), *T. b. brucei* GVR35 (Panel C), *T. b. rhodensiense UGA15* (Panel D) and *T. b. rhodensiense UGA17* (Panel E) were propagated in mice and processed (without permeabilization) for immunofluorescence analysis with anti-VSGm immune serum at dilutions of 1:4000 as described in the methods. The VSGm was detected in the various bloodstream form trypanosome species and strains.

#### Immunofluorescent staining of *T. b. brucei* in cattle brain histological sections

This was done on cattle brain sections similar to those in which H & E staining for trypanosome detection had previously been performed in another study. Signals were only observed in brain sections that had showed positive results with H & E staining but not the controls (sections which had showed negative results with H & E staining plus those which gave positive results but were probed with PBS or pre-immune serum) (Fig. 6). The experiment was repeated six times while yielding similar results.

**Fig. 6:**
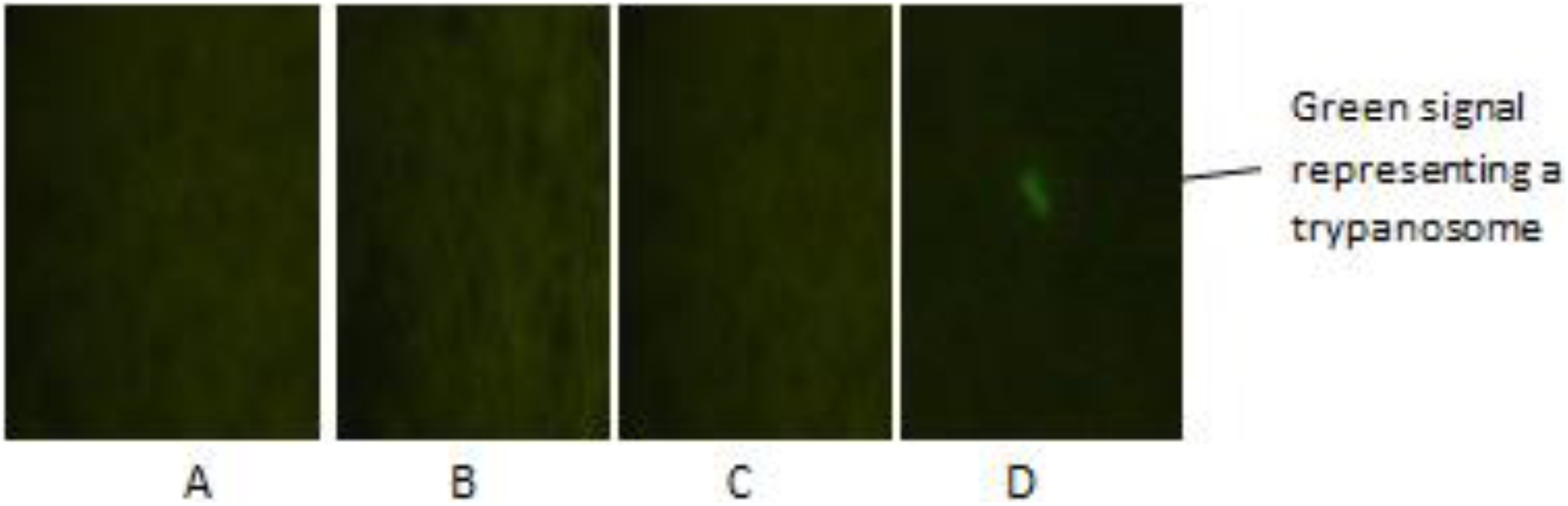
Immunofluoresent staining of *T. b. brucei* in infected cattle brain histological sections. Duplicate sections were first developed with H & E staining to ensure that trypanosomes were present before processing similar sections for immunofluorescence analysis. Section A had showed negative results with H & E staining and was probed with the immune serum and still showed no signals. Probed with pre-immune serum (Panel B), PBS (Panel C) and immune serum (Panel D). All primary antibodies were used at a dilution of 1:4000. A signal was only detected in Panel D.

## Discussion

This study has produced anti-VSG antibodies that have successfully detected trypanosomes in blood and cattle brain histological sections, and in addition, stained a broad range of bloodstream form pathogenic African trypanosome species and strains, exhibiting its potential to be used in detecting different species and strains of trypanosomes. In the preparation of VSGm, a protein of molecular weight approximately 61kDa that was observed was within the documented range of molecular weight of VSG monomer which is between 45 to 65kDa in *Trypanosoma brucei* and other closely related species, *T. evansi* and *T. equiperdum* (20). Actually, with the extensive recombination of genes and pseudo-genes, it is possible that the range may exceed the documented value. Xuedong Kang *et al* (21) reported the existence of a VSG molecule as small as 37kDa.

The presence of a single band of protein of approximately molecular weight 61kDa does not imply that only one variable antigen type (VAT) of VSGm was obtained because the primary difference between the VATs is in the sequence of amino acids in the highly hypervariable N-terminal domain of the VSG (22) consisting of about 350 to 400 residues which is approximately two thirds of the intact VSG molecule (23,24). If the extraordinarily conserved C-terminal GPI anchor signal sequence is disregarded, the range of sequence variation among VSGs is such that some sequences would be difficult to recognize as members of the family (22). The variation in sequence of amino acids would not cause a considerable difference in molecular weight thus the one band observed could represent several VATs. The findings that the antibody stained two strains of *Trypanosoma brucei [T. b. brucei* (427 and GVR35) and *T. b. rhodensiense (UGA 15 and UGA 17)]* plus *Trypanosoma congolense (BA59)* implies that more than just two VATs of VSGm could have been obtained.

It is not surprising that different VATs of VSGm could be obtained because it is known that at a given time, an individual bloodstream form trypanosome expresses only a single VSG gene, in a mutually exclusive fashion (25,26), from one of approximately 20 telomeric bloodstream form VSG expression site (ES) transcription units, which can be present at one or both ends of *T. brucei* chromosomes (25). There is thus a homogeneous display of identical surface epitopes in exposed amino (N)-terminal regions of the VSGm molecules (22). At regular intervals, a new VAT appears on the surface of the parasite, preventing recognition of the parasites by an upcoming anti-VSG response directed against the previous VAT (27). This therefore implies that it is possible to obtain different VATs when trypanosomes are propagated in various animals at different times.

Western blot analysis revealed a protein of similar molecular weight as the immunogen was recognized by the anti-VSG sera both in the bloodstream form trypanosome whole cell lysate and residue volumes of the immunogen. Neither trypanosome β-tubulin nor mammalian BSA were recognized implying that the sera were produced against the prepared protein that was used for immunization and this was confirmed by the observation that pre-immune sera at the same dilution did not give any signals. The finding that the *T. b. b.* VSGm immune sera produced showed a distinct uniform fluorescence of the entire morphology of bloodstream form trypanosomes but not procyclic form trypanosomes confirmed that the protein prepared was VSG because differentiation between the bloodstream and procyclic form trypanosomes is characterized by removal of the VSG surface coat and the acquisition of procyclin (28). It is the procyclin that dominates the procyclic cell surface (29).

Of great significance was the finding that the *T. b. b.* anti-VSGm immune sera produced distinct fluorescence of trypanosomes of other species (*T. congolense BA 59, T. b. rhodensiense UGA 15, T. b. rhodensiense UGA 17*) and strains (*T. b. brucei* GVR 35, *T. b. brucei* 427). This universal recognition could be via the C-terminal domain of VSG that is highly conserved at the homology region (30) or it could be the result of the multiple batch effect of producing enough variants found on a broad range of species/strains). Nevertheless, the observation is highly significant because it implies that the anti-VSGm produced can be used as a molecule to stain or locate more than one species of trypanosomes (with their various strains) *in vitro* especially in the diagnosis of trypanosomiasis. In addition to recognizing different strains of trypanosomes *in vitro*, the anti-VSGm immune serum produced a distinct fluorescence on the cattle brain tissue sections indicating the presence of trypanosomes in the brain sections. The study findings provide the possibility of using anti-VSGm immune serum for immunohistostaining of post mortem materials to localize trypanosomes in brain sections and other tissue sections while using another counterstaining reagent for the host cells. It can also be used as a diagnostic tool for the increased sensitive staining of trypanosomes in CSF and other body fluids to screen for the presence of trypanosomes both *in vitro* and *in situ*.

## Conclusions

The *T. b. brucei* anti-VSGm antibodies can be used detect various strains of *T. brucei* sub-species in tissues to monitor effectiveness of new drugs or vaccines in clearing the parasites or preventing their entry into the CNS. It can also be used to monitor the location of the parasites at the different stages of the disease and possibly to sensitively stain trypanosomes in CSF during staging of HAT. The anti-VSGm immune serum produced should be tested on more species and strains of trypanosomes or purified proteins to ascertain its effectiveness in staining different species/strains of trypanosomes to improve disease diagnosis.

## Authors’ contributions

**OR:** participated in study design, experimental and laboratory work, data analysis and manuscript preparation, **AP:** participated in study design, experimental and laboratory work, data analysis, **JN:** participated in experimental and laboratory work, data analysis and manuscript preparation, **NZ:** participated in study design, experimental and laboratory work, data analysis and manuscript preparation. **LGW:** conceived the project, participated in study design, data analysis and manuscript preparation. **NA**: participated in study design, experimental and laboratory work, data analysis and manuscript preparation.

## Acknowledgements

We are grateful to the staff at the COVAB, Makerere University, Small Animal Unit for the care and management of the experimental animals. We acknowledge the staff at the Molecular Biology Laboratory (MOBILA) at CoVAB, Makerere University for technical support in sample preparation and storage, laboratory work and data analysis. We also acknowledge Prof. Enock Matovu for providing African trypanosome stock reference strains, Prof. GW. Lubega for cattle brain histological sections, Ms. Monica Namayanja, Dr. Philip Magambo and Mr. Sylver Ochwo for guidance and useful discussions on immunization experimental and fluorescence laboratory work. This research work was fully self-funded.

